# Hormone therapy attenuates menopause-related impairments in motor cortical spike-timing-dependent plasticity

**DOI:** 10.64898/2026.11.12.765850

**Authors:** Paul Ansdell, Pádraig Spillane, Elisa Nédélec, Mollie O’Hanlon, Thomas Inns, Mathew Piasecki, Markus Hausmann, Stuart Goodall, Katy Vincent, Charlotte Stagg, Jessica Piasecki

## Abstract

The menopause marks the permanent cessation of menstruation via a decline in endogenous hormone production. This transition influences brain health, yet its effects on motor cortical excitability and spike-timing-dependent plasticity (STDP) are poorly understood.

Across two sites, 46 females were categorised as pre-menopausal (PRE, age 28±7 years), perimenopausal (PERI, 50±3 years), post-menopausal (POST, 60±8 years), or combined continuous hormone therapy users (HT, 52±3 years). Transcranial magnetic stimulation was used to measure corticospinal excitability (MEP/Mmax), short-interval intracortical inhibition (SICI), and intracortical facilitation (ICF) before, then 10 and 30 min after a paired associative stimulation (PAS) protocol (25-ms interstimulus interval). Serum hormone concentrations were quantified from venous blood.

Resting corticospinal excitability and SICI were not different between groups before the PAS protocol. However, a group × time interaction (p=0.0152) for MEP/Mmax revealed distinct STDP responses. PRE showed a significant increase in corticospinal excitability following PAS (+48%, p=0.0045), whereas PERI and POST showed no facilitation (p≥0.2079). Between groups, both PRE and HT exceeded PERI, and HT also exceeded POST; however, HT users showed a smaller, non-significant increase (+32%, p=0.0666) that collectively distinguished them from the blunted responses of PERI and POST. The magnitude of STDP was not associated with any single hormone or chronological age across the whole sample of participants.

These findings provide the first evidence that motor cortical STDP is influenced by menopause stage. With blunted responses in PERI and POST, yet preserved responses in HT users, these data have implications for motor learning and rehabilitation across the female adult lifespan.

## INTRODUCTION

The menopause demarcates the stage of a female’s life when menstruation permanently ceases due to the loss of ovarian reproductive function and is defined by 12 consecutive months without menstruation without using hormonal contraception (NICE, 2024). Prior to this point, females experience the peri-menopause transition, where reproductive hormone concentrations decline, with considerable variability (Grub et al., 2021). Throughout this transition, menopausal symptoms may negatively impact quality of life and are underpinned by physiological changes in a wide range of systems (Monteleone et al., 2018). Long term deteriorations in aspects of brain health, such as reduced brain glucose metabolism, reductions in grey and white matter volume, and increased amyloid-beta deposition have been previously observed following the menopause (Mosconi et al., 2021; Mosconi et al., 2018), which are collectively thought to increase the risk of neurodegenerative diseases. In addition, the menopause appears to act as an inflection point for accelerated neuromuscular deconditioning and plays a major role in the greater development of frailty in older females compared to males (Piasecki et al., 2024). Hormone therapy (HT) is often prescribed for the management of menopause-related symptoms and is generally considered to offer protective effects for the risk of neurodegenerative disease and cognitive decline (Kim et al., 2021; Maki et al., 2011). Despite this, the role of the menopausal transition and HT usage on neurophysiological function is less clear.

Key aspects of sensorimotor physiology, such as the ability to convert motor commands into forceful and/or accurate muscle contraction are impaired with age. Within the motor cortex, age-related reductions in _γ_-aminobutyric acid (GABA)-mediated inhibition are associated with poorer dexterity (Heise et al., 2013); while reductions in corticospinal tract excitability, have been observed in male or mixed-sex studies of ageing (Sale & Semmler, 2005; Škarabot, Ansdell, et al., 2019). As with much of the neuroscientific literature (Beery & Zucker, 2011), female-specific aspects of neurophysiological ageing are under-researched. In studies of pre-menopausal females, sex hormones have been demonstrated to alter cortical neurotransmission, with oestrogen potentiating intracortical facilitation, and progesterone, increasing inhibition (Ansdell et al., 2019; Smith et al., 2002; Smith et al., 1999). Additionally, Spillane et al. (2026) recently demonstrated how oestrogen increases excitability of the corticospinal tract. Specific data regarding how the stages of the menopause influence corticospinal and intracortical properties are lacking, with only Pesonen et al. (2021) demonstrating a longer TMS silent period in early perimenopause; however between-group differences in motor evoked potential (MEP) amplitude and a lack of MEP normalisation confound these findings. As such, a comprehensive assessment of intracortical neurotransmission and corticospinal excitability across stages of the menopause, with and without HT usage, is required to better understand how this hormonal transition might impact motor physiology.

One further sensorimotor function imperative to maintain into later life, is the ability to adapt in response to external or internal stimuli, termed neuroplasticity. Recent meta-analyses demonstrate an age-related reduction in neuroplasticity, as measured by the increase in MEP following a non-invasive brain stimulation paradigm (Shah et al., 2024). However, the authors highlighted considerable inter-individual variability in responses and discussed how the menopause, and sex hormones, must be either investigated or controlled in future research. Indeed, two studies have now reported that only females experienced an age-related decline in the response to paired associative stimulation (PAS), whereas males maintain the response into older age (Polimanti et al., 2016; Tecchio et al., 2008). Once more, the authors speculated that the menopause was a likely explanation of this finding, but did not directly study this in their investigations, with older groups aged ∼61 years and likely post-menopausal. Further support for the relationship between sex hormone concentrations and neuroplasticity can be found within literature concerning pre-menopausal females, with females only showing a facilitatory response to repetitive TMS on day 14 of the menstrual cycle (likely associated with high oestrogen concentrations), compared to a blunted response on day 1 (low oestrogen concentrations, Inghilleri et al. (2004). More recently, Spillane et al. (2026) demonstrated that the greatest response to PAS was observed in the mid-luteal phase of the menstrual cycle, although spike-timing-dependent plasticity (STDP) was observed in all phases. Evidence from ex vivo rodent models shows that oestrogen augments long-term potentiation (LTP) by enhancing N-methyl-D-aspartate (NMDA) receptor transmission (Smith & McMahon, 2005), yet a systematic investigation of how the various stages of the menopause, as well as the use of HT affect neuroplasticity in humans is lacking.

Accordingly, this study had two aims. Firstly, to investigate potential differences in corticospinal excitability and intracortical neurotransmission between groups of females from pre, to peri, and post-menopause, as well as HT users. Second, to investigate between group differences in the STDP response to PAS. We hypothesised that there would be a decrease in corticospinal excitability as sex hormones decreased from pre to post-menopause, which would also be accompanied by a decrease in the response to PAS. We also hypothesised that HT users would experience a greater degree of neuroplasticity compared to their counterparts.

## METHODS

### Ethical Approval

This study received institutional ethical approval from the Northumbria University Health and Wellbeing Research Ethics Committee (reference number: 39792) and Nottingham Trent University Invasive research ethics committee (reference number: 1606166) and was conducted according to the Declaration of Helsinki in all aspects apart from registration in a database. All participants gave their written informed consent prior to any part of the study.

### Sample Size Estimation

An a priori power calculation was performed using the effect size from Tecchio et al. (2008), who demonstrated an age-related decline in neuroplasticity in females, and reliability data from Ansdell et al. (2019). With parameters of α = 0.001, power = 0.99, ηp^2^ = 0.115, a minimum sample size of 48 participants between four experimental groups was calculated.

### Participants

Forty-six participants volunteered for the study and were categorised by an initial screening questionnaire as either pre-menopausal (PRE), peri-menopausal (PERI), post-menopausal (POST), or HT users according to the NICE Guidance on the Identification and Management of the Menopause (NICE, 2024). Specifically, pre-menopause (n = 12) was defined as a regular menstrual cycle length of 21-35 days, with no hormonal contraceptive usage or menstrual disturbance reported in the previous 12 months. Perimenopause (n = 12) was defined as individuals aged ≥40 years and reporting menstrual disturbance and menopausal symptoms, and were not using hormone therapies. Post-menopause (n = 12) was defined as individuals aged ≥40 years and ≥12 consecutive months without menstruation, and were not using hormone therapies. HT users (n = 10) were aged ≥40 years and used a combined, continuous exogenous HT (see details below). Demographic, anthropometric and hormonal data for all participants can be found in the results section. All participants were free from known neurological illness and musculoskeletal injury to the relevant limbs and were not taking any medication known to affect neurological function. Participants were instructed to refrain from strenuous physical activity and alcohol (24 h) and caffeine (12 h) before each experimental testing session. A multi-site approach was taken for this study, with n=29 participants tested at Northumbria University, and n=17 tested at Nottingham Trent University. All aspects of experimental design and methodologies described below, were identical at both sites, and researchers at each site were trained together prior to data collection to ensure methods were replicated appropriately.

### Experimental Design

This study was cross-sectional in design, with participants visiting the laboratory twice; once for familiarisation with experimental methods, then a subsequent experimental visit separated by at least 48 hours. Pre-menopausal participants completed the experimental visit between days 10-14 of their menstrual cycle to align with the late follicular phase and elevated oestrogen concentrations, which was confirmed post-hoc with serum hormone analyses (Spillane et al., 2026). HT users who were using transdermal patches completed the experimental visit within 72 hours of replacing their patch.

The familiarisation visit introduced participants to the procedures of brain and nerve stimulation, along with a full baseline assessment and a shortened (∼2 mins) PAS protocol. Experimental visits began with determination of stimulation intensity thresholds for electrical nerve stimulation, followed by TMS hotspot and resting motor threshold identification. Thereafter, a baseline assessment was completed and a PAS protocol performed. Assessments were then repeated 10- and 30-mins following PAS to measure the STDP response. Experimental visits finished with the collection of a venous blood sample for subsequent hormonal analyses. Detailed descriptions of each assessment are provided in the following sections.

### Procedures

#### Bipolar Surface Electromyography

Surface electromyography (EMG) was recorded from the first dorsal interosseous (FDI) muscle of the nondominant hand throughout assessments of nervous system function and PAS protocol. Two self-adhesive Ag/AgCl recording electrodes (20 × 15 mm, Neuroline 700, Ambu, Denmark) were placed ∼ 11 cm apart on the belly of the FDI, the reference electrode was placed over the ulnar styloid process of the same arm. The raw EMG signal was amplified (×1,000), band pass filtered (20–2,000 Hz), and digitised (5,000 Hz; 1401, CED, Cambridge, UK.

#### Electrical Stimulation

Activation of the FDI was achieved via percutaneous stimulation of the ulnar nerve, using surface electrodes (3.2 cm diameter; ValuTrode, Axelgaard Manufacturing, Denmark), with the cathode placed proximally to reduce the necessary current intensity (Pieber et al., 2015). The stimulating electrodes were connected to a constant current muscle stimulator (DS7AH, Digitimer, Welwyn Garden City, UK), which delivered single pulses of 200 _μ_s duration. During the delivery of electrical stimuli, participants were instructed to keep the target muscle relaxed, confirmed via visual inspection of the EMG trace. For Mmax assessment, stimulation intensity began at 30 mA and was increased by 10 mA until a plateau in peak-to-peak M-wave amplitude was observed. To ensure supramaximal stimulation and account for potential changes in axonal excitability, the intensity used for all Mmax assessments was increased by an additional 30%. Stimulation intensity (mean ± SD, PRE: 159 ± 20 mA, PERI: 144 ± 40 mA; HT: 130 ± 37 mA; and POST: 146 ± 40 mA; p = 0.3002) was not different between groups. For sensory stimulation as part of the PAS protocol, the same electrode placement and stimulus duration was utilised. Determination of perceptual threshold was achieved by starting at an intensity of 1 mA, which was increased in 1 mA increments until participants reported being able to feel the stimulus. This was confirmed by reducing the intensity by one increment, and re-testing whether the participants reported being able to feel the stimulus. To ensure effective sensory stimulation, the intensity was then increased to 300% of perceptual threshold (Stefan et al., 2000). Perceptual threshold was similar between groups (PRE: 4 ± 2 mA; PERI: 4 ± 1 mA; HT: 4 ± 1 mA; POST: 5 ± 1 mA; p = 0.3114).

#### Transcranial Magnetic Stimulation

Magnetic stimuli were delivered over the motor cortex contralateral to the nondominant limb, using a flat 70 mm figure of eight coil (Second Generation Remote 3190-00, Peak Magnetic Field 2.2 T), connected to two linked monopulse magnetic stimulators (Magstim BiStim^2^ and 200^2^, The Magstim Company, Whitland, UK). The TMS coil was orientated to induce a posterior-anterior directed magnetic field, delivering a 1-ms pulse. The “hotspot,” defined as the largest and most consistent motor-evoked potential (MEP) measured in the FDI, was located at the beginning of each visit, and the site was marked on the scalp with indelible marker pen to ensure consistent coil placement for subsequent stimulation. Resting motor threshold (rMT) was determined according to the IFCN definition (Rossini et al., 2015) as the lowest stimulation intensity (% Maximal Stimulator Output [MSO]) that induced a motor-evoked potential (MEP) in the relaxed FDI of >50 *µ*V in 5 of 10 trials. rMT was similar between groups (PRE: 54 ± 5%; PERI: 56 ± 13%; HT: 53 ± 8%; POST: 57 ± 9% MSO; p = 0.6809). This intensity was subsequently used to set the stimulation intensity for measuring corticospinal excitability, which was performed at 120% rMT. The peak-to-peak amplitude of MEPs (mV) was measured in Spike2 software (v10.08, CED, Cambridge, UK) and expressed as a % Mmax. Twenty unconditioned MEPs were recorded to collect a reliable within-session measure of corticospinal excitability. Visual inspection of each trial was performed, and any with voluntary activity in the preceding 100 ms were discarded.

Short-interval intracortical inhibition (SICI) was measured using a paired-pulse paradigm, with a conditioning pulse delivered at 80% rMT, followed by a test pulse delivered at 120% rMT, with an interstimulus interval of 2 ms (Kujirai et al., 1993). Intracortical facilitation (ICF) was measured using a paired-pulse paradigm, with a conditioning pulse delivered at 80% rMT, followed by a test pulse delivered at 120% rMT, with an interstimulus interval of 12 ms (Kujirai et al., 1993). For SICI and ICF, a total of 20 responses were recorded with each stimulation paradigm, with conditioned MEPs being expressed as a % of unconditioned MEPs. Due to a technical issue, paired-pulse TMS data was not present for n=2 PRE, n=3 PERI, and n=2 HT.

#### Paired Associative Stimulation

The PAS protocol consisted of 200 ulnar nerve stimuli, coupled with TMS, delivered at rest. Ulnar nerve stimulation was delivered at 300% perceptual threshold, and TMS was delivered at 120% rMT, with a 25 ms interstimulus interval, aligning with the majority of PAS studies (Carson & Kennedy, 2013). Paired stimuli were delivered at a frequency of 0.25 Hz, except for instances where participants had an rMT ≥60% (n=3 PRE, n=3 PERI, n=1 HT, and n=6 POST), where frequency was reduced to 0.2 Hz allowing the TMS unit time to recharge between stimuli. Sensitivity analyses were therefore run to assess whether the different frequency of PAS affected results. For the MEP/Mmax linear mixed effects model, rMT (≥60 vs <60) was first included as a covariate, then models were run on participants with rMT <60 only. In both instances, the interaction F statistic was consistent (F = 2.81 to 2.32) and significant (all p < 0.05) indicating that PAS frequency had no influence on STDP. The magnitude and time course of STDP were assessed by repeated assessments of MEP/Mmax, SICI, and ICF at 10- and 30-mins after the PAS protocol (Wischnewski & Schutter, 2016).

#### Blood Sampling and Hormone Analysis

Venous blood samples were collected at the end of each testing session to avoid any potential impact that stress and subsequent increased cortisol levels might have on the STDP response to PAS (Sale et al., 2008). Each blood draw consisted of a 10 mL sample collected from an antecubital vein, which was left upright for 2 h to coagulate in clot activator vacutainer tubes. Samples were centrifuged for 15 min at 1,000 *g*, and the serum supernatant was then pipetted into aliquots and stored at −80°C until hormone analysis. Hormone analyses were performed at the Bioanalytical Facility, University of East Anglia (Norwich, UK) and undertaken in Good Clinical and Laboratory Practice conditions. Serum concentrations of testosterone (nmol.L-1), androstenedione (nmol.L-1), 17-hydroxyprogesterone (nmol.L-1), dehydroepiandrosterone (nmol.L-1), and cortisol (nmol.L-1) were measured by liquid chromatography-tandem mass spectrometry (LC-MS/MS) on a Waters Xevo TQ-XS coupled with an Acquity I-class system (Waters Corp., Milford, MA, USA), with each analyte quantified by targeted multiple reaction monitoring. Steroids were extracted from 100 µL of serum, with deuterated internal standards, by supported liquid extraction (ISOLUTE SLE+, Biotage, Uppsala, Sweden) and eluted with methyl tert-butyl ether. Assays were calibrated using NIST SRM971-traceable serum-based calibrators (MassChrom Steroids Panel 1 & 2, Chromsystems, Munich, Germany), with multilevel quality controls (MassCheck, Chromsystems) analysed in each run. Inter- and intra-assay coefficients of variation (CV) were 3.5–8.6% across their respective assay ranges, and the lower limit of quantification was 0.1 nmol.L-1. Serum 17β-oestradiol (pmol.L-1) and progesterone (nmol.L-1) were measured by electro-chemiluminescence immunoassay on the COBAS e601 automated platform (Roche Diagnostics, Mannheim, Germany), with inter-assay CVs of ≤3% within their respective analytical working ranges. The minimum detectable concentrations were 18.4 pmol.L-1 and 0.2 nmol.L-1 for 17β-oestradiol and progesterone, respectively. Blood samples were not obtained for 1 premenopausal, 4 perimenopausal, 2 HT users, and 4 post-menopausal participants; therefore, comparisons between groups contain 11 PRE, 8 PERI, 8 HT, and 8 POST.

#### Statistical Analysis

All data processing and analyses were conducted using R Statistical Software (v4.6.1). Alpha level was set at 0.05 for all tests, and all data were visualised in ggplot (Wickham, 2016). For measures with multiple observations at each timepoint (e.g., MEP/Mmax), data are presented as estimated marginal means ± 95% confidence intervals from model outputs. Linear mixed effects models (lme4, Bates et al. (2015); lmerTest, Kuznetsova et al. (2017) for Satterthwaite-approximated degrees of freedom) were used to determine whether variables of interest could be predicted by the fixed effects of group (PRE, PERI, HT and POST), and time (baseline, +10 mins, +30 mins following PAS), as well as group x time interactions. Random effects included intercepts for participant and measurement block (time) within participants. To assess differences at baseline, the model structure was “outcome ∼ group + (1|participant)”; whereas to assess STDP following PAS, the model structure was “outcome ∼ group x time + (1|participant/time)”. Models were fitted by residual maximum likelihood (REML) estimation. Outcomes (Mmax, MEP/Mmax, SICI, and ICF) were natural-log transformed prior to modelling to satisfy residual normality, and estimates were back-transformed for reporting (geometric means, and contrasts expressed as ratios). Estimated marginal means and pairwise contrasts were obtained with emmeans (Lenth & Piaskowski, 2026). Between-group differences at each timepoint were assessed using Tukey’s adjustment for all pairwise comparisons; within-group changes from baseline (+10 min and +30 min versus baseline) were assessed using Dunnett’s adjustment for comparisons against a single reference level.

For measures with a single observation at each timepoint or trial (e.g., hormone concentrations), data are presented as mean ± standard deviation for normally distributed data or geometric mean ± confidence intervals for data that required logarithmic transformation. One-way analyses of variance (ANOVA) were used to determine differences between groups. The normality of distribution was confirmed using Shapiro-Wilk tests. Hormone concentrations were log transformed prior to the ANOVA to ensure normal distribution, then back transformed for presentation. Where a main effect of group was detected, post hoc pairwise comparisons were performed with Tukey’s adjustment.

Exploratory Spearman rank correlations were performed between the change in MEP/Mmax at 30 minutes post-PAS (calculated as the ratio of per-participant geometric means at 30 minutes to baseline) and each of the hormone concentrations across the whole sample.

## RESULTS

### Participant characteristics and hormonal status

Demographic, anthropometric, and hormonal data for each group are presented in Table 1 and Figure 1. As anticipated, PRE were younger than all other groups (F_3,42_ = 71.103, p < 0.0001), with POST older than PERI (p = 0.0006) and HT (p = 0.0131). There were no between group differences in stature or body mass (p ≥ 0.0947).

**Figure 1:**
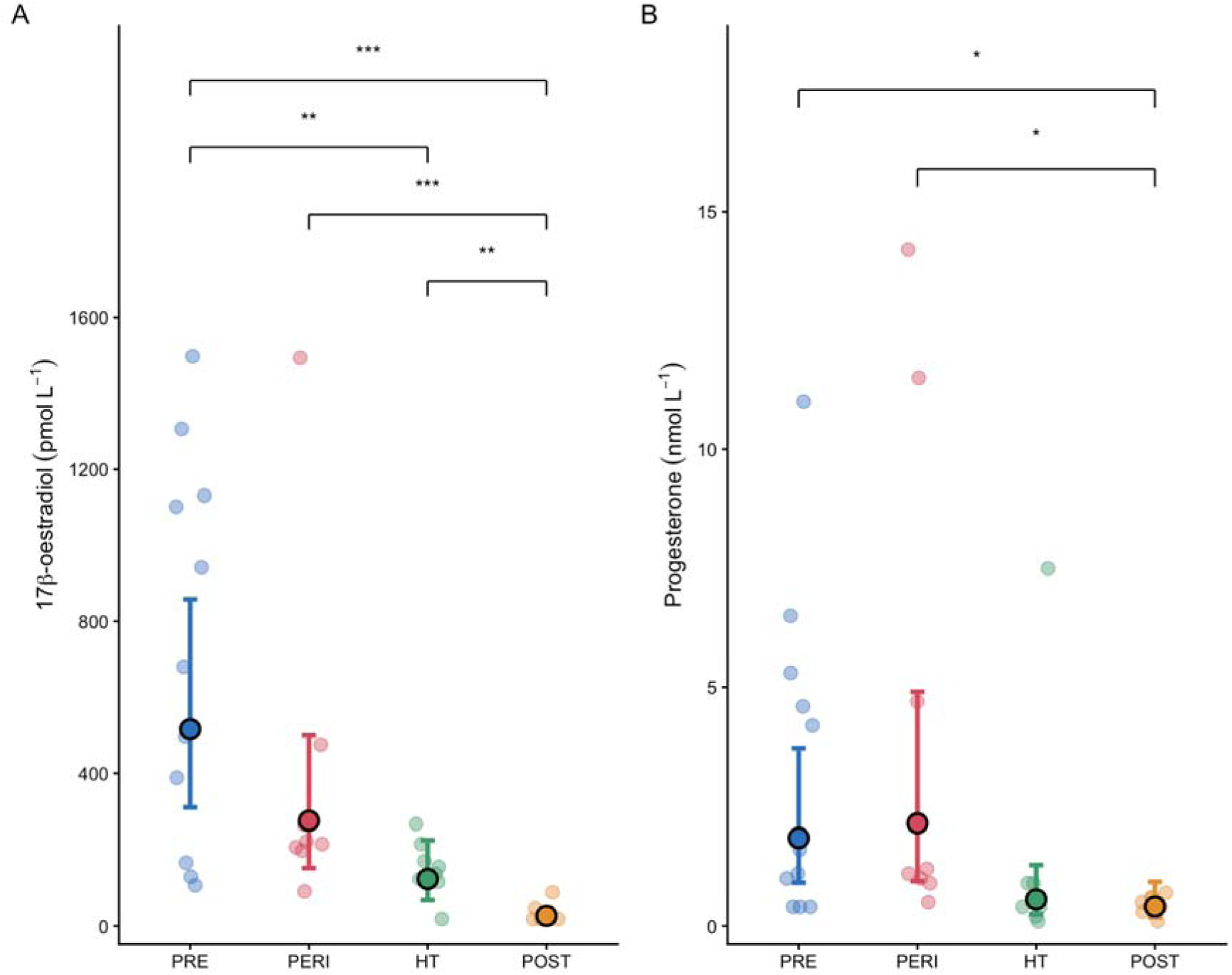
Sex hormone concentrations of the four groups; 17_β_-Oestradiol (Panel A) and Progesterone (Panel B). PRE = pre-menopausal females (n=11); PERI = perimenopausal females (n=8); HT = hormone therapy users (n=8); POST = post-menopausal females (n=8). Individual participants are represented by the small dots, with bold dots and error bars representing the back-transformed geometric mean and 95% confidence intervals following logarithmic transformation. * p < 0.05; ** p < 0.01; *** p < 0.001.

**Table 1:** Age and anthropometric data from pre-menopausal, perimenopausal, HT users, and post-menopausal participants. Data are presented as mean ± standard deviation.

|  | **Pre-menopause**<br>**(n=12)** | **Perimenopause**<br>**(n=12)** | **HT Users**<br>**(n=10)** | **Post-menopause**<br>**(n=12)** |
|---|---|---|---|---|
| Age (years) | 28 ± 7 | 50 ± 3 | 52 ± 3 | 60 ± 8 |
| Stature (cm) | 165 ± 4 | 163 ± 6 | 168 ± 4 | 165 ± 4 |
| Body mass (kg) | 66.5 ± 8.7 | 67.5 ± 10.2 | 69.2 ± 7.0 | 70.1 ± 7.7 |

Hormonal profiles for each group are presented in Table 2, with the anticipated differences in ovarian hormones 17β-oestradiol (F_3,31_ = 21.377, p < 0.0001) and progesterone (F_3,31_ = 4.564, p = 0.0092, Figure 1). Post hoc contrasts demonstrated that all groups had greater 17β-oestradiol concentrations than POST (p ≤ 0.0041), and that PRE had greater concentrations than HT (p = 0.0042). For progesterone, PRE and PERI were greater than POST (p ≤ 0.0375). Further between group differences are highlighted within Table 2.

**Table 2:**
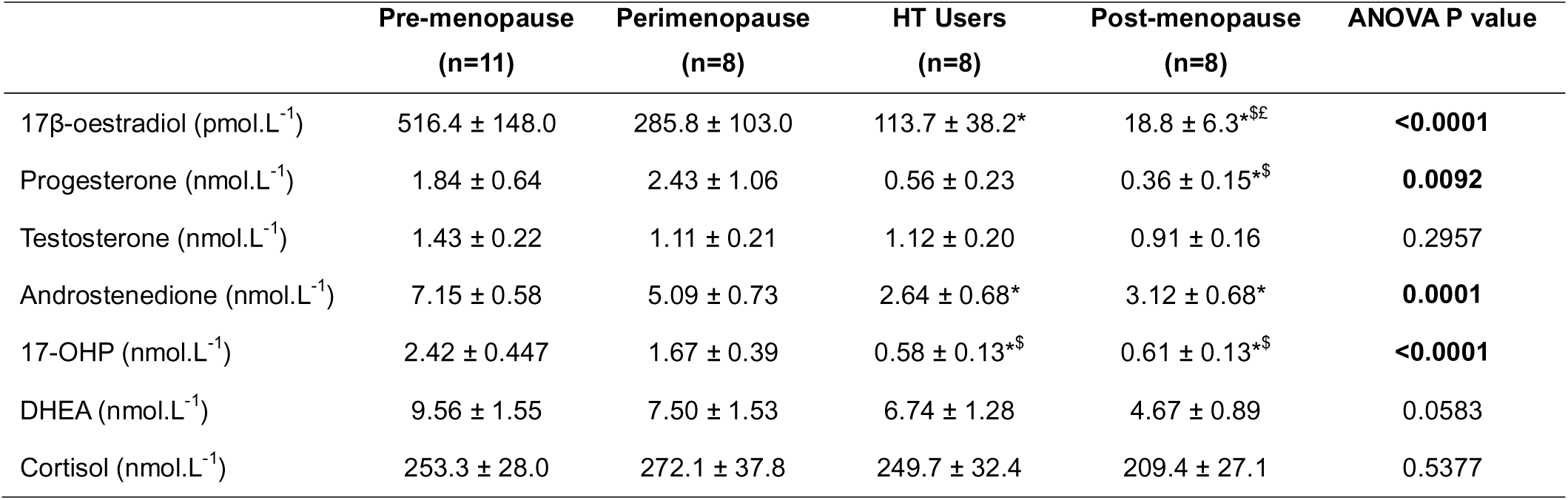
Hormone concentrations from the blood samples analysed from pre-menopausal, perimenopausal, HT users, and post-menopausal participants. Data are presented as geometric mean ± standard error. * = less than premenopause, $ = less than perimenopause, £ = less than hormone therapy users. 17-OHP = 17-Hydroxyprogesterone; DHEA = dehydroepiandrosterone; ANOVA = one-way analysis of variance.

|  | **Pre-menopause<br>(n=11)** | **Perimenopause<br>(n=8)** | **HT Users<br>(n=8)** | **Post-menopause<br>(n=8)** | **ANOVA P value** |
|---|---|---|---|---|---|
| 17β-oestradiol (pmol.L<sup>-1</sup>) | 516.4 ± 148.0 | 285.8 ± 103.0 | 113.7 ± 38.2* | 18.8 ± 6.3*<sup>\$£</sup> | **<0.0001** |
| Progesterone (nmol.L<sup>-1</sup>) | 1.84 ± 0.64 | 2.43 ± 1.06 | 0.56 ± 0.23 | 0.36 ± 0.15*<sup>\$</sup> | **0.0092** |
| Testosterone (nmol.L<sup>-1</sup>) | 1.43 ± 0.22 | 1.11 ± 0.21 | 1.12 ± 0.20 | 0.91 ± 0.16 | 0.2957 |
| Androstenedione (nmol.L<sup>-1</sup>) | 7.15 ± 0.58 | 5.09 ± 0.73 | 2.64 ± 0.68* | 3.12 ± 0.68* | **0.0001** |
| 17-OHP (nmol.L<sup>-1</sup>) | 2.42 ± 0.447 | 1.67 ± 0.39 | 0.58 ± 0.13*<sup>\$</sup> | 0.61 ± 0.13*<sup>\$</sup> | **<0.0001** |
| DHEA (nmol.L<sup>-1</sup>) | 9.56 ± 1.55 | 7.50 ± 1.53 | 6.74 ± 1.28 | 4.67 ± 0.89 | 0.0583 |
| Cortisol (nmol.L<sup>-1</sup>) | 253.3 ± 28.0 | 272.1 ± 37.8 | 249.7 ± 32.4 | 209.4 ± 27.1 | 0.5377 |

Of the HT users, 9 used transdermal oestradiol patches or gels (0.289 ± 0.346 mg/day), including one participant using a combination of transdermal gel and pessaries (0.010 mg/day), and one participant who used pessaries alone (0.010 mg/day). All participants used daily oral progestogens, with either medroxyprogesterone acetate (n=2, 3 mg/day) or micronised progesterone (n=7 100 mg/day, n=1 200 mg/day) being used.

### Spike-timing-dependent plasticity following paired associative stimulation

The Mmax evoked was not different between groups (F_3,42_ = 2.681, p = 0.0590), and there was no effect of time (F_2,84_ = 1.687, p = 0.1913). At baseline specifically, there were no differences between groups (PRE = 10.09 [7.78, 13.09]; PERI = 10.59 [8.16, 13.74]; HT = 9.29 [6.99, 12.35]; POST = 6.60 [5.09, 8.57] mV; p ≥ 0.0796).

There was a main effect of group (F_3,42.008_ = 4.611, p = 0.0071) for MEP/Mmax, but there was no effect of time (F_2,85.210_ = 0.948, p = 0.3916). There was, however, a group x time interaction effect (Figure 2, F_6,85.215_ = 2.811, p = 0.0152). The baseline-only model indicated no effect of group before the PAS intervention (F_3,42.224_ = 1.865, p = 0.1502), but differences between groups emerged 10 mins after the PAS protocol, with PRE and HT having larger MEP/Mmax than PERI (p ≤ 0.0185). Then at 30 mins, both PRE and HT were again greater than PERI (p ≤ 0.0311), whilst HT were also greater than POST (p = 0.0301). Indeed, within-group comparisons indicated that PRE experienced significant increases at 10 mins (48 [11, 91] % increase, p = 0.0045) and 30 mins (28 [1, 73] % increase, p = 0.0418). HT users did not demonstrate an increase at 10 mins (p = 0.6094) and despite a 32 [-2, 78] % increase, did not reach significance at 30 mins (p = 0.0666). PERI and POST did not demonstrate any within-group changes (p ≥ 0.2079). There were no significant correlations between individual hormone concentrations and the change in MEP/Mmax at 30 minutes for the full sample (all p ≥ 0.5162, n = 35).

**Figure 2:**
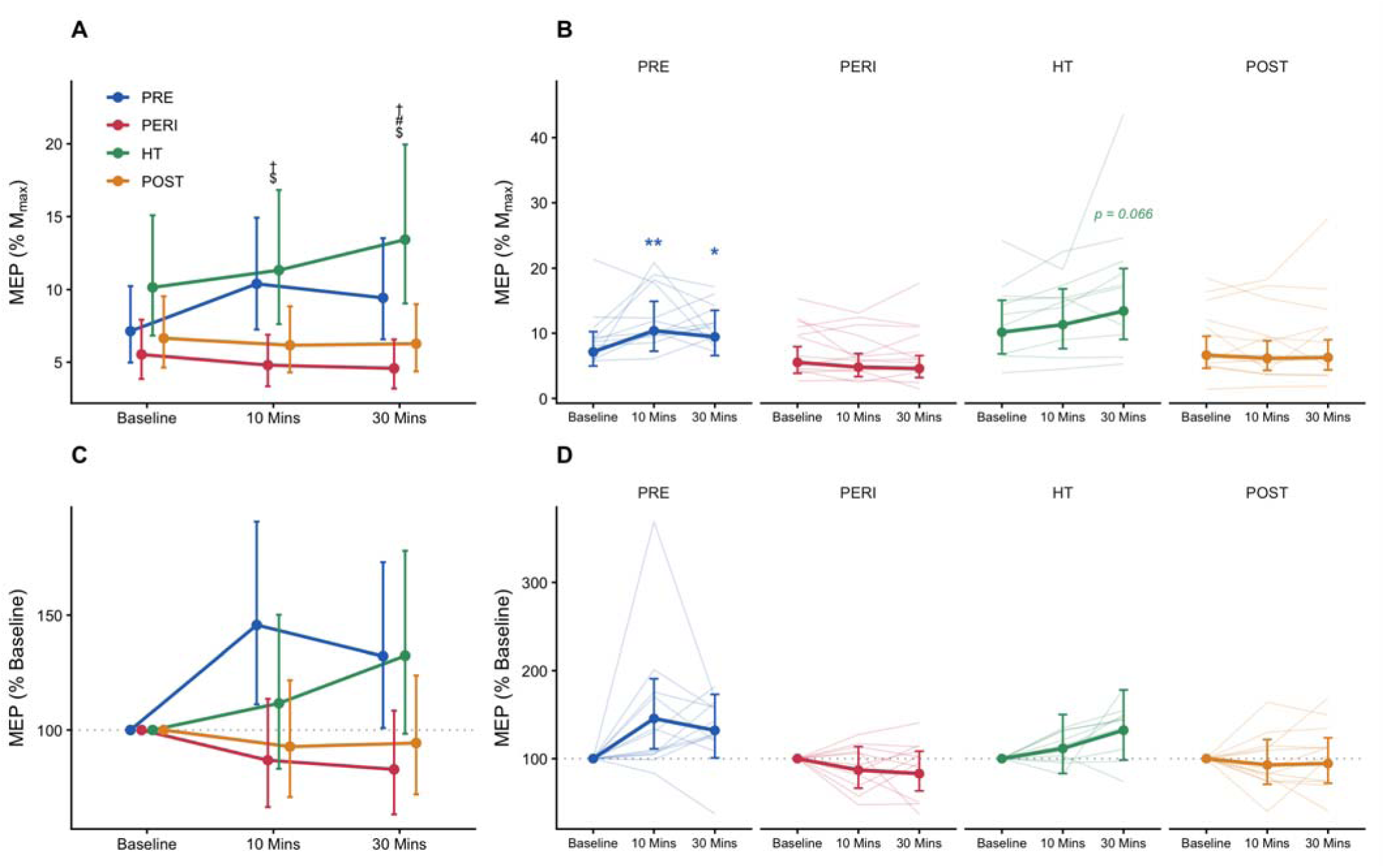
Motor evoked potential (MEP) data before and after the paired associative stimulation (PAS) protocol. Panel A displays the transformed estimated marginal means and confidence intervals for each group’s MEP normalised to M wave at each oint. Panel B displays individual participant responses (thin lines) as well as the same estimated marginal means and dence intervals as Panel A. Panel C and D are similar, but present data as % baseline to aid within group comparisons. No ntial statistics were run on the MEP data normalised to % baseline. PRE = pre-menopausal females (n=12); PERI = enopausal females (n=12); HT = hormone therapy users (n=10); POST = post-menopausal females (n=12). † = PRE greater PERI (p < 0.05), $ = HT greater than PERI, # = HT greater than POST, * = greater than baseline (p < 0.05), ** = greater than ine (p < 0.01).

Two sensitivity analyses were run to further explore this group x time interaction. Firstly, adding participant age as a covariate left the group x time interaction unchanged (p = 0.015), with no independent age effect (p = 0.889) or age x time interaction (p = 0.861). Secondly, a model that excluded the two HT participants using synthetic progestins (medroxyprogesterone acetate) retained the group x time interaction effect (p = 0.009), however the 30 mins timepoint was significantly greater than baseline for the HT group (42 [2, 98] % increase, p = 0.034).

For SICI (Figure 3), there were no group (F_3,35.099_ = 1.292, p = 0.2924), time (F_2,70.238_ = 0.463, p = 0.6315), or group x time interaction effects (F_6,70.234_ = 0.659, p = 0.6828). Indeed, the baseline-only model confirmed that there were no differences between groups before the PAS intervention (F_3,35.124_= 1.920, p = 0.1448).

**Figure 3:**
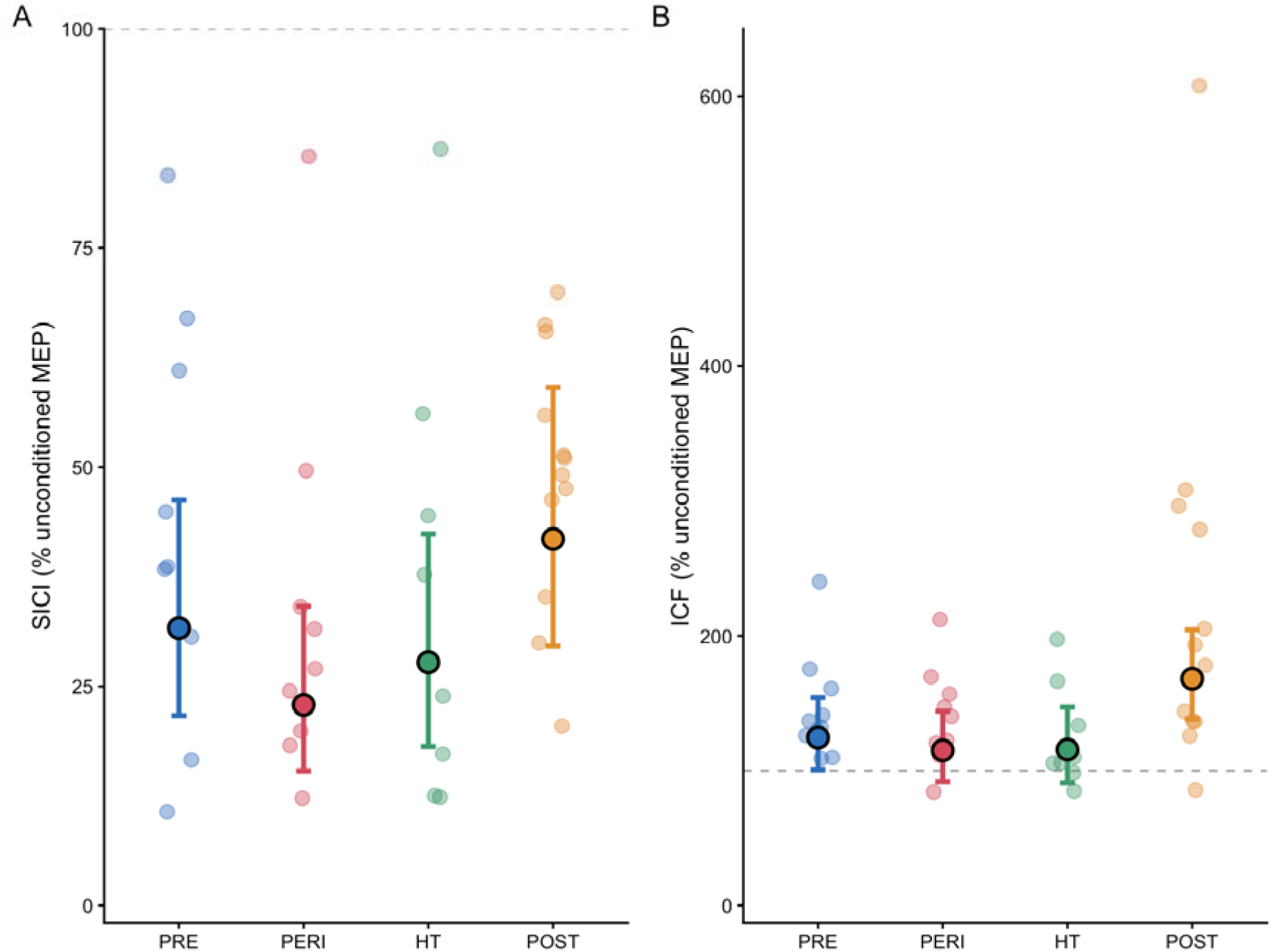
Data from paired pulse transcranial magnetic stimulation (TMS) obtained before the paired associative stimulation protocol (PAS). Panel A displays individual data, as well as the back-transformed estimated marginal means and confidence intervals for short-interval cortical inhibition (SICI), and Panel B displays the same data for intracortical facilitation (ICF). PRE = pre-menopausal females (n=10); PERI = perimenopausal females (n=9); HT = hormone therapy users (n=8); POST = post-menopausal females (n=12).

For ICF (Figure 3), there were no group (F_3,35.275_ = 1.962, p = 0.1375), time (F_2,68.777_ = 0.435, p = 0.6494), or group x time interaction effects (F_6,68.779_ = 1.732, p = 0.1266). The baseline-only model demonstrated a main effect of group (F_3,35.414_ = 3.146, p = 0.0370), however, there were no significant post-hoc contrasts, despite POST (168 [138–205] % unconditioned MEP) having greater geometric mean values compared to PERI (115 [92–144] % unconditioned MEP, p = 0.0643) and HT (116 [91-147] % unconditioned MEP, p = 0.0849) before the PAS intervention.

## DISCUSSION

This study demonstrates that menopausal status and exogenous hormone therapy usage affect the degree of spike-timing-dependent plasticity that females experience following a paired associative stimulation intervention. While corticospinal excitability and intracortical inhibition were not different before the intervention, differences between groups emerged following PAS, with corticospinal excitability in PRE (+48%) and HT (+32%) exceeding that of PERI and POST, who had entirely blunted responses. These data support recent observations reporting that changes in sex hormone concentrations across the menstrual cycle impact neuroplasticity (Spillane et al., 2026). Here, changes in neuroplasticity were not associated with changes in a single hormone or chronological age across the whole sample; instead, the reduction in the combined hormonal milieu following the menopause, and its partial replacement with HT, influences the dynamic mechanisms of neuroplasticity within the motor cortex across the adult female lifespan.

### Spike-timing-dependent plasticity across the female lifespan

The primary finding of this study was that the magnitude of STDP following PAS was blunted in PERI and POST, compared to PRE who experienced a facilitatory response. Alongside this, HT users, who were of similar age to PERI, experienced a facilitatory response that was more akin to PRE. While this study is the first to categorise participants by menopausal status, sex differences in the age-related response to PAS have previously been reported (Polimanti et al., 2016; Tecchio et al., 2008). For instance, Polimanti et al. (2016) concluded that older females (∼61 years) had a blunted response compared to their younger counterparts (∼29 years), and the degree of PAS-induced plasticity was positively correlated with concentrations of testosterone. The present study supports the former conclusion, with the average age of PRE (28 years) and POST (60 years) being similar to those previously reported (Polimanti et al., 2016). However, the present study extends these findings into midlife and highlights that perimenopause appears to be a critical period within the female lifespan for the maintenance of neuroplasticity. This age-related decrease in cortical neuroplasticity in response to interventions such as PAS is well established, with older adults (typically >60 years) often demonstrating reduced responses compared to younger adults (Müller-Dahlhaus et al., 2008). A recent systematic review on this topic highlighted how this age-related effect was accompanied by substantial heterogeneity within and between studies (Shah et al., 2024), with previous work demonstrating that protocol and neurophysiological differences between groups do not explain this heterogeneity (Sale et al., 2007). Accordingly, this study advances the collective understanding of age-related changes in motor cortical plasticity, providing evidence menopausal status, appears to underpin this age-related adaptation.

Other evidence for the influence of sex hormones on neuroplasticity comes from pre-menopausal females, and repeated measures studies across the menstrual cycle. These cyclical changes differ from the menopause, where changes are permanent, but offer an alternative model to study their effects. Recently, our group observed that the response to PAS was greatest in the mid-luteal phase, where concentrations of oestrogens and progesterone are elevated (Spillane et al., 2026). This supports the present study’s findings where PRE experienced a greater magnitude of plasticity. One additional factor to consider is that PRE were tested mid-cycle in the present study, rather than the mid-luteal phase. Given the potential mechanisms by which oestrogen and progesterone might interact with STDP, as discussed below, this could imply that the age-related decrease in PAS response is likely to be even greater if the mid-luteal phase is used as the reference point. Contrary to the present study, Spillane et al. demonstrated that plasticity was able to be induced even in menstrual cycle phases associated with low sex hormone concentrations, whereas POST in the present study did not experience plasticity. Other investigations that have used interventions such as repetitive TMS also observed an entirely blunted response in the early follicular (low hormone) phase of the menstrual cycle (Inghilleri et al., 2004), whereas intermittent-theta-burst stimulation appeared to be potentiated in the follicular phase (Ramdeo et al., 2024), implying that hormonal effects are likely task-specific. Despite this, studies that employ PAS to elicit spike-timing-dependent plasticity across the female lifespan agree that timepoints or groups with low sex hormone concentrations experience a blunted response to the protocol, compared to high-hormone conditions (Polimanti et al., 2016; Spillane et al., 2026; Tecchio et al., 2008).

### Shared mechanisms between endogenous and exogenous hormones

Another novel aspect of the present study is that HT users experienced a greater magnitude of STDP compared to PERI, who were of a similar age (52 vs. 50 years). In vivo human evidence into the neurophysiological effects of exogenous hormones is comparatively sparse, compared to endogenous hormonal effects, but the potential mechanisms are somewhat likely shared. The HT users in the present study primarily used a combination of transdermal oestradiol and an oral progestogen. Oestradiol increases NMDA receptor density (Woolley et al., 1997) and augments long-term potentiation (Smith & McMahon, 2005), which provides a common mechanism for the greater STDP in the PRE and HT groups. Transdermal oestrogen might also facilitate STDP through the serotonergic system; in postmenopausal women, Moses et al. (2000) demonstrated that a combination of oestradiol and micronized progesterone (similar to the HT regimes used in this study), increased cortical 5-HT2A receptor availability. Pharmacological increases in serotonergic tone (i.e., SSRIs) augment the facilitatory response to PAS at a 25 ms ISI, while abolishing the inhibitory response at a 10 ms ISI (Batsikadze et al., 2013). While the serotonergic route is relatively speculative, the oestradiol-related potentiation of PAS is likely an explanation for the similar responses in PRE and HT.

The progestogen contribution to STDP is less clear compared to the effect of oestradiol’s. Spillane et al. (2026) observed the greatest PAS response in pre-menopausal females during the mid-luteal phase, when endogenous progesterone is elevated. The authors suggested that in this phase, progesterone and its metabolite allopregnanolone promote NMDA-induced dopamine release (Cabrera & Navarro, 1996; Rougé-Pont et al., 2002), which would potentiate responses to PAS (Kuo et al., 2008). On the other hand, the mid-luteal phase is also a high oestrogen phase, unlike mid-cycle where pre-menopausal females were tested in the present study, which could suggest a facilitatory effect of combined hormone elevation. Whereas other evidence suggests that progestogens applied alone (e.g., medroxyprogesterone acetate) reduced dopamine availability (Tandon et al., 1983). However, it is important to note that the progesterone concentrations observed in the present study were low (all groups <3 nmol.L^-1^) due to testing taking place mid-cycle, in comparison to values observed in the mid-luteal phase (∼40 nmol.L^-1^, Spillane et al. (2026)). This means that progesterone’s mechanistic role in STDP across the female lifespan is less clear than that of oestrogen. However, this does provide rationale and highlights the gap in contemporary understanding about the way in which progesterone and progestogens might influence the mechanisms of motor cortical neuroplasticity in humans. Collectively, the similar STDP response in PRE and HT groups in contrast with PERI and POST, in the present study is likely underpinned by shared mechanisms related to oestradiol, and although the source might differ (i.e., endogenous vs. exogenous), the neuroactive effects are similar.

### No differences in resting excitability

The absence of any between group differences in corticospinal excitability and SICI contradicts our hypothesis, as well as work in premenopausal women across the menstrual cycle. In previous work, MEP/Mmax peaked when oestrogen concentrations are greatest in the late follicular phase (Spillane et al., 2026), and SICI was greatest in the mid-luteal phase (Ansdell et al., 2019; Smith et al., 2002). Given the large between-group differences in 17β-oestradiol observed in the present study (Table 2), parallel differences in excitability might have been anticipated as well; however, there are several factors that might reconcile these differences. Firstly, the cyclical modulation of MEP/Mmax in pre-menopausal women results in a modest within-participant effect (∼2% Mmax, Spillane et al. (2026)) from a repeated measures study design. In contrast, the models within the present cross-sectional study had approximately half the variance in the MEP/Mmax explained by participant and timepoint (see supplementary material), highlighting the known variability of TMS-derived measures at rest (Darling et al., 2006). There was, however, a main effect of group in ICF at baseline, but the post hoc pairwise comparisons were insignificant when corrected for multiple comparisons (Figure 3). The geometric means suggest that POST had greater ICF, and less SICI than other groups, but the confidence intervals around the point estimate were wide. Collectively, these factors imply that a repeated measures study offers a more sensitive way of quantifying the effects of sex hormones on corticospinal excitability directly, which is somewhat less feasible when studying the menopause compared to the menstrual cycle.

Another cross-sectional study across the menopause observed greater MEP amplitudes in early perimenopausal compared to late perimenopausal females, during muscle contraction (Pesonen et al., 2021). The authors suggested that this decrease was related to menopause-related changes in intracortical inhibition, as MEPs were correlated with the corticospinal silent period duration. However, given that MEPs were not normalised to Mmax, which represents a confounding factor in cross-sectional investigations of corticospinal excitability (Škarabot et al., 2021), caution should be taken when interpreting these data. Additionally, the relationship between MEP amplitude and silent period duration is unlikely to represent any physiological link between the two variables, as this is a known methodological link that has led researchers to suggest a CSP:MEP ratio is more likely to uncover mechanistic links between the two outcomes (Orth & Rothwell, 2004; Škarabot, Mesquita, et al., 2019). Given the methodological considerations of the two outcomes, these between group differences should be considered with caution.

### Further considerations

A strength of the study was that all HT users were using combined, continuous hormone therapy, minimising the cyclical hormonal variation that occurs in the PRE group. But while dosages and delivery methods were reported by participants, the methods employed did not capture or fully standardise the administration or timing of HT usage before experimental visits. These factors influence the pharmacokinetics of transdermal hormones and oral progestogens, and how they are detected in serum. For example, oral micronized progesterone concentrations peak within 2-3 hours of consumption (Simon et al., 1993), and with this often being taken before sleep, it is likely that the daytime testing in the present study missed peak concentrations, despite a half-life of ∼17 hours (Simon et al., 1993). Additionally, participants in the HT group were a mixture of those using micronized progesterone (n=8) or medroxyprogesterone acetate (n=2), with the latter being known to bind to more receptor types than just progesterone alone. The sensitivity analysis for MEP/Mmax that excluded the two participants using medroxyprogesterone acetate revealed a post-hoc difference from baseline at the 30 mins time point for the HT group, whereas this was not significant with them included. This should be caveated with the fact that the p values were almost equidistant from the threshold of significance (p = 0.0666 vs. p = 0.0341), and firm conclusions about the different neurophysiological effects of synthetic progestins vs. bio-identical progesterone should not be drawn from this analysis. However, this does highlight an avenue for future research to explore; and whether the effects of different hormone therapies, and their timings and dosages, impact neurophysiological outcomes should be investigated.

This study attributed menopause-related differences in STDP to mechanisms associated with the changes in sex hormone concentrations across this hormonal transition, but it is also established that PAS-induced plasticity is state-dependent, with factors such as sleep deprivation (Kuhn et al., 2016), chronic stress (Gellner et al., 2022), and prior physical activity (Mang et al., 2014) influencing outcomes. These factors were not measured in the present study yet represent potential ways in which the menopause could influence STDP as well as direct neurophysiological mechanisms. Conceivably, these factors would fit with the between-group differences observed in the present study, with PERI typically being the most symptomatic group experiencing the weakest response to PAS. Likewise, HT users might have experienced symptom relief, which could contribute to their facilitatory response. This possibility is speculative yet complementary to the direct hormonal hypothesis tested in this study, but it emphasises the value of characterising menopause-related symptom burden alongside neurophysiological outcomes in future work.

Finally, although the present study studied the PRE group between days 10-14 of their menstrual cycle in order to test participants in a high oestrogen state. Based on the progesterone data in Figure 1, it appears that one participant met the criteria to be considered within the luteal phase (>9.5 nmol.L^-1^, Chinta *et al.,* 2020). Likewise, two of the PERI group also had progesterone values over this threshold. Considering this, and as the timing of ovulation was not verified with methods such as urinary luteinising hormone tests, this manuscript refers to the time of testing for PRE as ‘mid-cycle’ rather than the late follicular phase.

## CONCLUSION

This study provides the first evidence that spike-timing-dependent plasticity of the motor cortex is affected by menopausal status. Despite comparable resting excitability between groups, the plasticity response to paired associative stimulation was blunted in perimenopausal and post-menopausal females yet preserved in hormone therapy users, whose response resembles that of pre-menopausal females. Given that these conclusions were still retained after including age as a covariate, this implies a hormonal mechanism underpins the neurophysiological differences. However, despite between group differences in hormone concentrations, the differences in plasticity were not attributable to any single hormone, which implicates the broader changes to the hormonal milieu as a determinant of motor cortical plasticity across the female lifespan. Spike-timing-dependent plasticity within the motor cortex underpins a wide range of motor learning principles, therefore these neurophysiological findings carry direct consequences for how motor learning and rehabilitation are approached around the menopausal transition. The findings also make the case that the menopause is a fundamental biological principle that should not be overlooked or ignored, and it represents a critical window in the female lifespan that has long-term consequences for healthy ageing.

## Acknowledgements

The authors thank Dr Rachel Dunn and Dr Jonathan Tang for the analysis of the blood samples.

## Funding

This was supported by a Royal Society Research Grant (JP and PA, RGS/R1/231080) and a Physiological Society Research and Knowledge Exchange Award (PA and CS).

## Notes

### Competing Interest Statement

The authors have declared no competing interest.

