## Supplementary material for "Hormone therapy attenuates menopause-related impairments in motor cortical spike-timing-dependent plasticity"

**Supplementary Material: Linear mixed effects model fit**

For MEP/Mmax (n = 46), the model explained 54% of the variance in the log-transformed data (conditional R²), and the fixed effects of group, time and their interaction alone explained 13% (marginal R²). The adjusted ICC indicated that 47% of the variance not explained by the fixed effects came from differences between participants (SD = 0.55) and between measurement blocks within participants (SD = 0.25). The remainder reflected stimulus-to-stimulus residual variability (SD = 0.65; all SDs on the natural-log scale). This degree of clustering is consistent with the known between-participant and between-block variability of TMS-evoked responses, and supports the nested random-intercept structure.

For SICI (n = 39), the model explained 57% of the variance in the log-transformed data (conditional R²), with a marginal R² of 5%. The adjusted ICC indicated that 55% of the variance not explained by the fixed effects came from differences between participants (SD = 0.55) and between measurement blocks within participants (SD = 0.29), with the remainder reflecting stimulus-to-stimulus residual variability (SD = 0.56). Like MEP/Mmax, this clustering supports the nested random-intercept structure.

For ICF (n = 39), the model explained 24% of the variance in the log-transformed data (conditional R²), with a marginal R² of 4%. The adjusted ICC indicated that 21% of the variance not explained by the fixed effects came from differences between participants (SD = 0.20) and between measurement blocks within participants (SD = 0.19). Most of the unexplained variance therefore reflected stimulus-to-stimulus residual variability (SD = 0.54). Clustering was weaker than for MEP/Mmax and SICI, but the nested random-intercept structure was kept because it reflects the repeated-measures design and keeps the models consistent across outcomes.
